# Identification of a PRC2 accessory subunit required for subtelomeric H3K27 methylation in Neurospora

**DOI:** 10.1101/2020.01.06.896845

**Authors:** Kevin J. McNaught, Elizabeth T. Wiles, Eric U. Selker

## Abstract

Polycomb repressive complex 2 (PRC2) catalyzes methylation of histone H3 lysine 27 (H3K27) in genomic regions of most eukaryotes and is critical for maintenance of the associated transcriptional repression. However, the mechanisms that shape the distribution of H3K27 methylation, such as recruitment of PRC2 to chromatin and/or stimulation of PRC2 activity, are unclear. Here, using a forward genetic approach in the model organism *Neurospora crassa*, we identified two alleles of a gene, *NCU04278*, encoding an unknown **<u>P</u>**RC2 **<u>a</u>**ccessory **<u>s</u>**ubunit (PAS). Loss of PAS resulted in losses of H3K27 methylation concentrated near the chromosome ends and derepression of a subset of associated subtelomeric genes. Immunoprecipitation followed by mass spectrometry confirmed reciprocal interactions between PAS and known PRC2 subunits, and sequence similarity searches demonstrated that PAS is not unique to *N. crassa*. PAS homologs likely influence the distribution of H3K27 methylation, and underlying gene repression, in a variety of fungal lineages.

## INTRODUCTION

Specification and maintenance of cellular fate in higher eukaryotes requires the concerted effort of transcription factor networks and chromatin-modifying complexes. Polycomb repressive complex 2 (PRC2), which catalyzes methylation of lysine 27 on histone H3 (H3K27), is one such chromatin-modifying complex essential for normal development in many organisms (1, 2). The three core components of PRC2, which are conserved in plants, animals, and fungi (3, 4), are EED, SUZ12, and the methyltransferase EZH2. Human PRC2 additionally associates with RBBP4/7, as well as other accessory proteins, which define the two human PRC2 subtypes: PRC2.1 and PRC2.2 (5). Analogous subtypes of PRC2 have also been described in *Drosophila melanogaster* (6–8) while in *Arabidopsis thaliana*, distinct PRC2 complexes are defined by different paralogs of the core PRC2 member, SUZ12 (9). Here, using a genetic approach, we provide evidence that a previously unidentified accessory subunit of PRC2 is responsible for functional specialization in the filamentous fungus *Neurospora crassa*.

Neurospora bears homologs of the human core PRC2 components EED (EED), SUZ12 (SUZ12), and EZH2 (SET-7), as well as RBBP4/7 (NPF) (10). While loss of either EED, SUZ12, or SET-7 completely abolishes H3K27 methylation, which normally covers ∼7% of the genome (10), loss of NPF affects H3K27 methylation in a region-specific manner (10). Considering that homologs of NPF are components of other chromatin-modifying complexes (11), it remained possible that the H3K27 methylation defect in Δ*npf* strains is not entirely attributable to PRC2 dysfunction. It is of obvious interest to identify all protein players involved in the establishment and maintenance of H3K27 methylation and delineate their respective roles. Using a forward genetic selection for factors defective in Polycomb silencing in *N. crassa* we report the isolation of mutant alleles of a previously uncharacterized gene (*NCU04278*) necessary for subtelomeric H3K27 methylation and silencing of associated genes. Immunoprecipitation followed by mass spectrometry of NCU04278-interacting proteins demonstrated that NCU04278 is a **<u>P</u>**RC2 **<u>a</u>**cccessory **<u>s</u>**ubunit, and we therefore named it PAS. PAS homologs are present in lineages of both Sordariomycetes and Leotiomycetes, suggesting that PAS may play a crucial role in regulating PRC2 in a wide variety of fungal species.

## RESULTS

### Isolation, mapping and identification of potential Polycomb group gene, *pas (NCU04278)*

We recently described a genetic scheme to select for mutants defective in Polycomb silencing in *N. crassa* (12). Briefly, a wild-type strain bearing two antibiotic-resistance genes repressed by H3K27 methylation, *hph* and *nat-1*, is subjected to ultraviolet radiation and mutants resistant to both Hygromycin B and Nourseothricin are isolated. One mutant recovered in this fashion, which we now designate *pas^UV1^*, led to antibiotic-resistance comparable to loss of the EZH2 homolog, SET-7, which is responsible for all known H3K27 methylation in *N. crassa* (Fig. 1A) (10). We mapped the causative mutation in the *pas^UV1^* strain, which was in the Oak Ridge genetic background, by crossing it to the highly polymorphic “Mauriceville” wild-type strain (13), pooling genomic DNA from antibiotic-resistant progeny, and scoring the percentage of Oak Ridge single nucleotide polymorphisms (SNPs) from whole-genome sequencing data (14). Oak Ridge SNPs were enriched on the right arm of linkage group (LG) V in a region that included a frameshift mutation in *NCU04278* (M660fs; <u>A</u>TG -> <u>TT</u>TG) (Fig. 1B).

**FIG 1.**
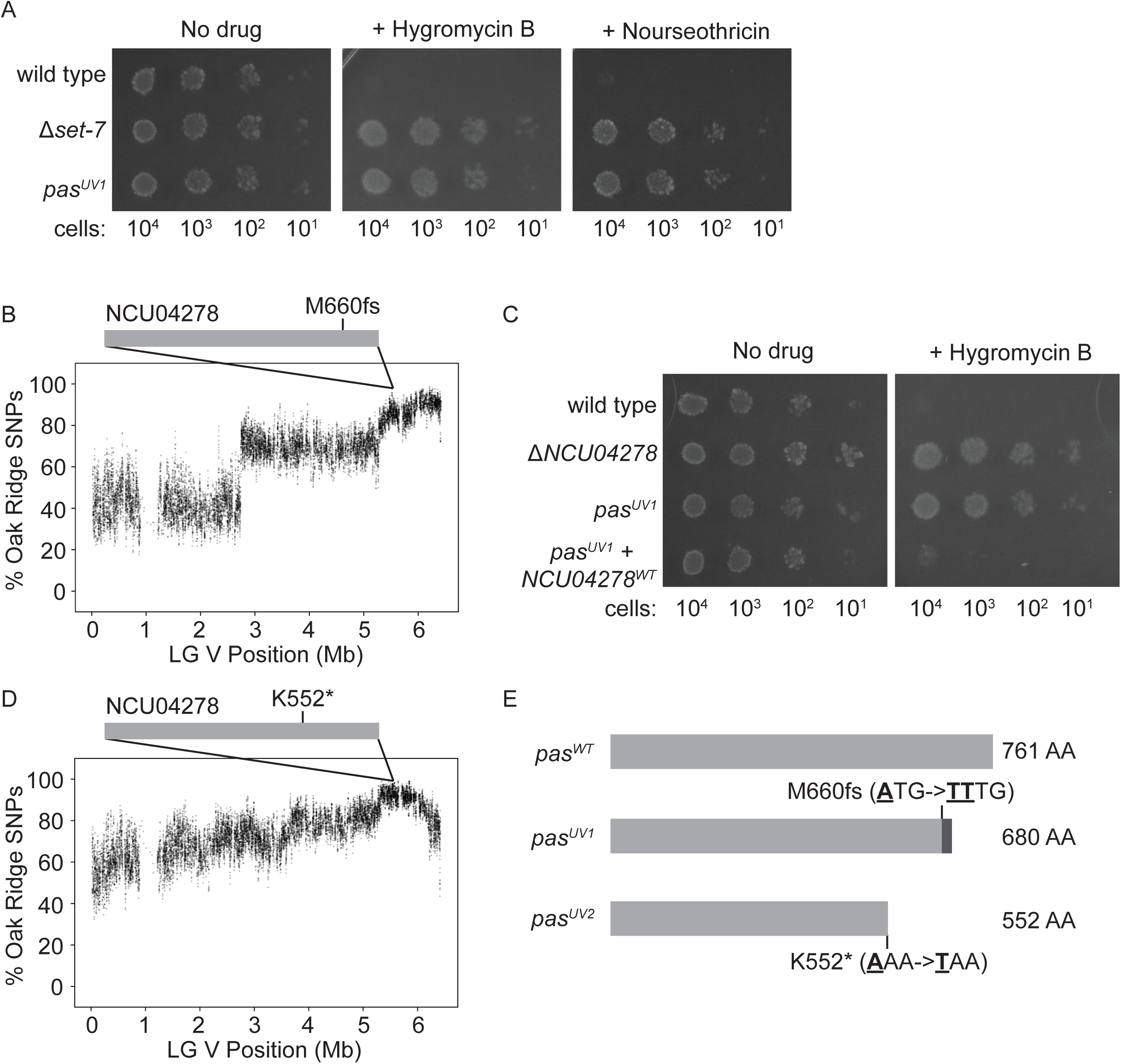
Identification of *NCU04278* as a potential component of the Polycomb silencing pathway. (A) Serial dilution spot test silencing assay for the indicated strains plated on the indicated media. All strains harbor *PNCU05173::hph* and *PNCU07152::nat-1*. (B) Whole genome sequencing of pooled *pas^UV1^* mutant genomic DNA identified a region on the right arm of LG V which is enriched for Oak Ridge single nucleotide polymorphisms (SNPs) and contained an insertional frameshift in *NCU04278* (protein structure is represented as a solid gray bar, as it contains no predicted domains). Each point represents a running average (window size = 10 SNPs, step size = 1 SNP). (C) Serial dilution spot test silencing assay for the indicated strains. *pas^UV1^* + *NCU04278^WT^* has a wild-type copy of *NCU04278* at the *his-3* locus. All strains harbor *PNCU05173::hph*. (D) As in B, but for *pas^UV2^* which contains a premature stop codon in *NCU04278*. (E) Schematic of the primary sequence and structural changes of *NCU04278* in the two *pas* alleles.

To verify that the observed mutation in *NCU04278* is the pertinent mutation in *pas^UV1^*, we targeted a wild-type copy of *NCU04278* to the *his-3* locus in a *pas^UV1^* mutant strain. The ectopic copy of *NCU04278* complemented the *pas^UV1^* mutant*, i.e*., restored Hygromycin B sensitivity (Fig. 1C). In addition, deletion of the wild-type allele of *NCU04278* was sufficient to confer Hygromycin B resistance (Fig. 1C). We subsequently isolated and mapped a second allele of *NCU04278* (*pas^UV2^*; K552*; <u>A</u>AA -> <u>T</u>AA) (Fig. 1D,E), further confirming the involvement of *NCU04278* in the antibiotic-resistant phenotype.

### PAS is necessary for silencing subtelomeric H3K27-methylated genes

Although our forward genetic selection was designed to identify novel components of the Polycomb repression pathway, in principle, mutations might confer resistance to Hygromycin B and Nourseothricin in some manner independent of derepression of the H3K27-methylated antibiotic-resistance genes, such as by stimulating drug efflux (15) or by a global effect on transcription, leading to non-specific derepression of *hph* and *nat-1*. To determine if loss of PAS has specific defects in Polycomb silencing, we performed mRNA-seq on Δ*pas* and wild-type strains in biological replicate and compared the gene expression profiles with previously generated wild-type and Δ*set-7* data sets (16). We found that 33 genes were upregulated, and 12 downregulated, greater than two-fold in Δ*pas* strains compared to wild-type strains (Fig. 2A,B). Although less than 9% of all genes are H3K27-methylated in a wild-type strain, 64% of the upregulated genes in Δ*pas* strains were in this select group. Moreover, there was significant overlap between the upregulated genes in Δ*pas* and Δ*set-7* strains (*P* = 6.095 x 10^-29^), although loss of SET-7 appeared to derepress more H3K27-methylated genes (Fig. 2A), *i*.*e*., Δ*pas* strains derepress a subset of SET-7 targets.

**FIG 2.**
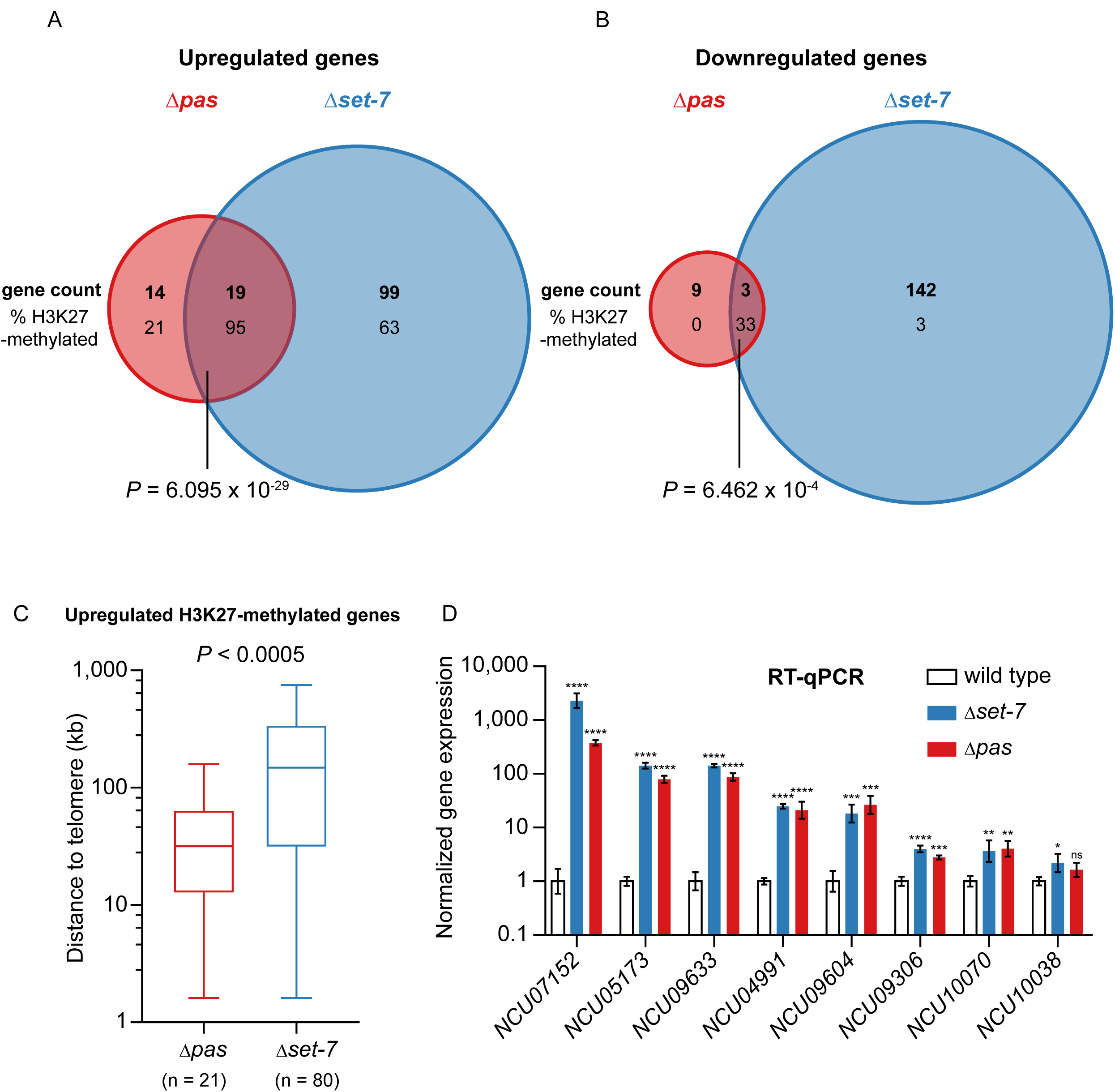
Loss of PAS upregulates a subset of SET-7-repressed genes near chromosome ends. (A) Venn diagram depicting genes (numbers in bold) that appear upregulated by mRNA-seq in both *Δpas* and *Δset-7* strains, only in *Δpas* strains, or only in *Δset-7* strains, using a significance cutoff of log2(mutant/wild type) > 1 and *P* value < 0.05. The percentage of total upregulated genes that are H3K27-methylated for each gene set is indicated below the total gene count. Significance of overlapping gene sets was determined using a hypergeometric test. (B) As in A, but for downregulated genes (log2(mutant/wild type) < -1). (C) Box and whisker plot of the distances from the telomere of H3K27-methylated genes that appear upregulated by mRNA-seq in the indicated genotypes. Boxes represent interquartile range, horizontal lines represent the median, and whiskers represent minimum and maximum values (*P* < 0.0005, two-tailed Mann-Whitney test). (D) RT-qPCR of H3K27-methylated genes that were replaced with antibiotic resistance genes (*NCU07152* and *NCU05173*) and used for initial selection of mutants, and H3K27-methylated genes that appeared upregulated in both Δ*pas* and Δ*set-7* strains by mRNA-seq (*NCU09633*, *NCU04991*, *NCU09604*, *NCU09306*, *NCU10070*, *NCU10038*). Each value was normalized to expression of *NCU02840* (39) and is presented relative to wild type. Filled bars represent the mean from biological triplicates and error bars show SD (**** for *P* < 0.0001, *** for *P* < 0.001, ** for *P* < 0.01, * for *P* < 0.05, and ns for not significant; all relative to wild type by two-tailed, unpaired t-test).

We previously demonstrated that there are at least two categories of H3K27 methylation in *N. crassa*: telomere-dependent and telomere-independent (17). Because loss of PAS only affected a subset of SET-7 targets, we considered the possibility that PAS may be responsible for specifically silencing one category of H3K27-methylated genes, *i*.*e*., those that are telomere-proximal or telomere-distal. We therefore analyzed the genomic location of H3K27-methylated genes upregulated in Δ*pas* and Δ*set-7* strains, respectively, and found that the genes derepressed by loss of PAS are closer to the chromosome ends than are the genes derepressed by loss of SET-7 (*P* < 0.0005) (Fig. 2C). This suggested that PAS is responsible for silencing telomere-proximal H3K27-methylated genes.

We verified the results of our mRNA-seq experiments by performing reverse transcription followed by quantitative polymerase chain reaction (RT-qPCR) on RNA isolated from biological triplicates of wild-type, Δ*set-7*, and Δ*pas* strains (Fig. 2D). In addition to confirming the derepression of the native genes replaced by the antibiotic-resistance genes in the selection strain (*NCU07152* and *NCU05173*), we confirmed a significant increase in gene expression for five out of six H3K27-methylated genes that appeared upregulated by mRNA-seq in both Δ*pas* and Δ*set-7* strains (Fig. 2D). We conclude that PAS represses a subset of SET-7 targets that are telomere-proximal.

### PAS is necessary for subtelomeric H3K27 methylation

Considering that the loss of PAS led to the derepression of H3K27-methylated genes near the chromosome ends, we wondered if there was also a concomitant loss of H3K27 methylation at subtelomeric regions. To address this possibility, we performed H3K27me2/3 chromatin immunoprecipitation followed by sequencing (ChIP-seq) on two Δ*pas* siblings and compared the results with the distribution of H3K27me2/3 in a wild-type strain (18) (Fig. 3A). We found that the general distribution of H3K27me2/3 in Δ*pas* strains was comparable to wild-type strains, except for clear losses near the chromosome ends (Fig. 3A). A western blot for H3K27me3 confirmed that Δ*pas* strains had reduced global H3K27me3 (Fig. 3B). Detailed analysis of the H3K27me2/3 ChIP-seq revealed that 284 genes in Δ*pas* strains had at least two-fold reductions in H3K27me2/3 compared to wild-type strains (blue dots; Fig. 3C) and 34 genes showed at least two-fold greater H3K27me2/3 than wild-type strains (red dots; Fig. 3C). To confirm the H3K27me2/3 ChIP-seq results, we performed H3K27me2/3 ChIP followed by qPCR (ChIP-qPCR) on wild-type, Δ*set-7*, and Δ*pas* strains in biological triplicate. We confirmed the findings for three regions expected to have no change in H3K27me2/3 (*NCU05086*, *NCU08085*, and *NCU08251*), three regions with expected losses in H3K27me2/3 (*NCU05173*, *NCU07152*, and Telomere IIIR), and one region with an expected gain in H3K27me2/3 (*NCU07801*) (Fig. 3D). Thus, PAS is critical for normal subtelomeric H3K27 methylation.

**FIG 3.**
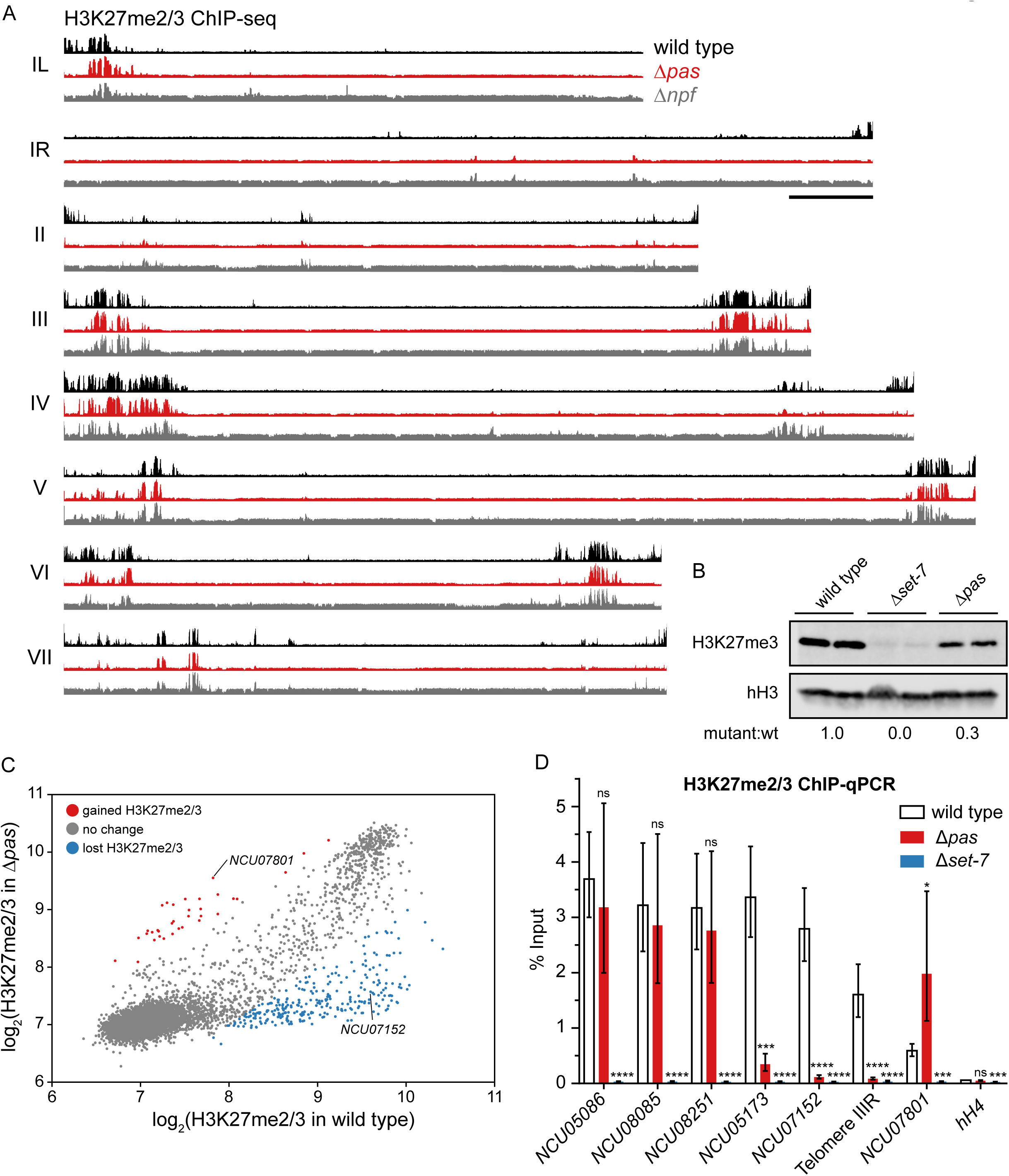
Loss of PAS results in global reduction of H3K27 methylation. (A) H3K27me2/3 ChIP-seq tracks from merged biological replicates of wild-type (black), *Δpas* (red), and *Δnpf* (gray) strains displayed for all seven Linkage Groups (LGs) of *N. crassa*. LG I is split at the right end of its centromere into IL and IR. Y-axes are 900, 850, and 450 RPKM for wild type, *Δpas*, and *Δnpf* respectively. Black bar represents 500 kb. (B) Western blot showing H3K27me3 and total histone H3 (hH3) in the indicated strains. Biological replicates are shown. The same lysate was analyzed on separate gels, and hH3 was used as a sample processing control. The number displayed below each genotype is the average ratio of mutant H3K27me3 level (normalized to hH3 level) to wild-type H3K27me3 level (normalized to hH3 level). (C) Scatter plot showing the correlation of H3K27me2/3 levels at all genes (dots) in wild type and *Δpas* based on merged biological replicates of ChIP-seq data. Dots representing genes are colored based on their relative H3K27me2/3 level in *Δpas* compared to wild type (red for log2(*Δpas*/wild type) > 1, gray for -1 < log2(*Δpas*/wild type) < 1, and blue for log2(*Δpas*/wild type) < -1). Representative genes that gained (*NCU07801*) or lost (*NCU07152*) H3K27me2/3 in *pas* are indicated. (D) H3K27me2/3 ChIP-qPCR to validate ChIP-seq data at eight regions – *NCU05086*, *NCU08085*, *NCU08251* (retained H3K27me2/3); *NCU05173*, *NCU07152,* Telomere IIIR (lost H3K27me2/3 in *Δpas*); *NCU07801* (gained H3K27me2/3 in *Δpas*); *hH4* (negative control). Filled bars represent the mean of biological triplicates and error bars show SD (**** for *P* < 0.0001, *** for *P* < 0.001, * for *P* < 0.05, ns for not significant; all relative to wild type by two-tailed, unpaired t-test).

### Strains lacking PAS have H3K27 methylation defects similar to Δ*npf* strains

Loss of subtelomeric H3K27 methylation has been previously observed in strains lacking the PRC2 component, NPF (10). For this reason, we repeated H3K27me2/3 ChIP-seq on Δ*npf* siblings to compare with the distribution of H3K27me2/3 observed in Δ*pas* strains (Fig. 3A). We found that while the distribution of H3K27 methylation in each strain was unique, the overall changes in the two strains were similar (Fig 4A). Indeed, comparison of H3K27me2/3 levels over all genes using Spearman’s correlation coefficient showed that Δ*npf* and Δ*pas* strains are more similar to each other than they are to wild type strains (Fig. 4B). Interestingly, unlike Δ*npf* strains (10), strains lacking PAS do not exhibit a linear growth defect (Fig. 4C). Therefore, the phenotypic consequences of losing NPF or PAS are not equivalent.

**FIG 4.**
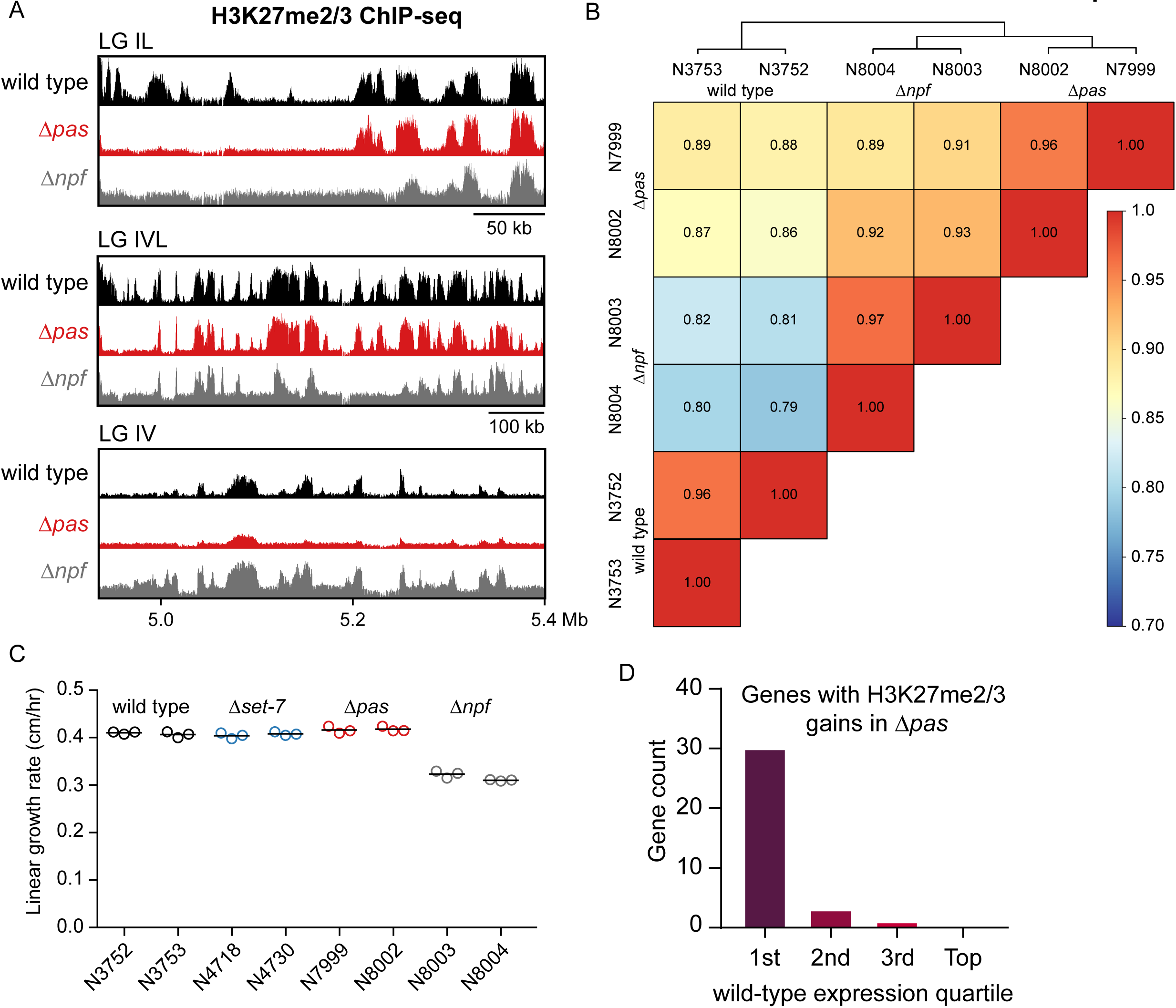
H3K27 methylation defects in *Δpas* and *Δnpf* are similar. (A) H3K27me2/3 ChIP-seq of merged biological replicates of wild-type (black), *Δpas* (red), and *Δnpf* (gray) strains for select genomic regions. Y-axes are 1000, 1000, and 500 RPKM for wild type, *Δpas*, and *Δnpf* respectively. (B) Spearman correlation matrix comparing H3K27me2/3 levels at all genes in ChIP-seq biological replicates of wild-type, *Δpas*, and *Δnpf* strains. Spearman correlation coefficients are displayed, and associated boxes are color-coded according to the legend. (C) Linear growth rates measured by ‘race tubes’ (40) are shown for two biological replicates of wild-type (N3752, N3753), *Δset-7* (N4718, N4730), *Δpas* (N7999, N8002), and *Δnpf* (N8003, N8004) strains. Horizontal lines represent the mean of three technical replicates (open circles). (D) Analysis of wild-type gene expression level of genes that gain H3K27me2/3 in *Δpas* strains.

### Relationship between gene expression changes and H3K27 methylation levels in Δ*pas* strains

To test for a correlation between the gene expression changes in Δ*pas* strains and the losses or gains of H3K27 methylation in Δ*pas*, we examined the intersection of these two data sets (see supplemental material). We found that while the loss of H3K27 methylation in Δ*pas* strains was not sufficient for gene activation, all but one (*NCU08790*) of the 21 H3K27-methylated genes upregulated in Δ*pas* strains lost H3K27 methylation (see supplemental material). In contrast, there was no overlap between the genes that gained H3K27 methylation in Δ*pas* strains and the genes that were downregulated Δ*pas* strains (see supplemental material). Analysis of the genes that gained H3K27me2/3 in Δ*pas* strains revealed that the majority (88%) fall into the lowest quartile of gene expression in wild-type strains (Fig. 4D). In summary, the losses of H3K27 methylation in Δ*pas* strains are associated with, but not sufficient for, increased gene transcription, and the gains of H3K27 methylation in Δ*pas* strains are mostly on genes that are normally lowly transcribed.

### PAS is an accessory subunit of PRC2

To determine if PAS acts in a protein complex, we immunopurified PAS-3xFLAG from *N. crassa* lysate and identified its copurifying proteins by mass spectrometry. In addition to PAS itself, we identified the four known components of PRC2: SET-7, SUZ12, EED, and NPF (Fig. 5A) (10). We also immunopurified SUZ12-3xFLAG and a negative control (3xFLAG-EPR-1) (12), and analyzed their copurifying proteins by mass spectrometry. In addition, we re-examined previously collected mass spectrometry data from a 3xFLAG-EED purification (10). The purifications of the known PRC2 components, SUZ12 and EED, yielded PAS as well as the rest of the known members of PRC2 (Fig. 5A). The negative control (3xFLAG-EPR-1) purification did identify peptides of NPF but did not detect any peptides of SET-7, SUZ12, EED, or PAS (Fig. 5A). Besides the proteins listed in Fig. 5A, the only other proteins with peptides detected in the PAS, SUZ12, and EED purifications, but absent in the control (EPR-1) were NCU01249 (importin α), NCU02407 (dihydrolipoyl dehydrogenase), and NCU06482 (pyruvate dehydrogenase). Thus, it appears that PRC2 in *N. crassa* can form a five-member complex and that PAS is an accessory component. For this reason, we have designated NCU04278 a **<u>P</u>**RC2 **<u>a</u>**ccessory **<u>s</u>**ubunit (PAS) (Fig. 5B).

**FIG 5.**
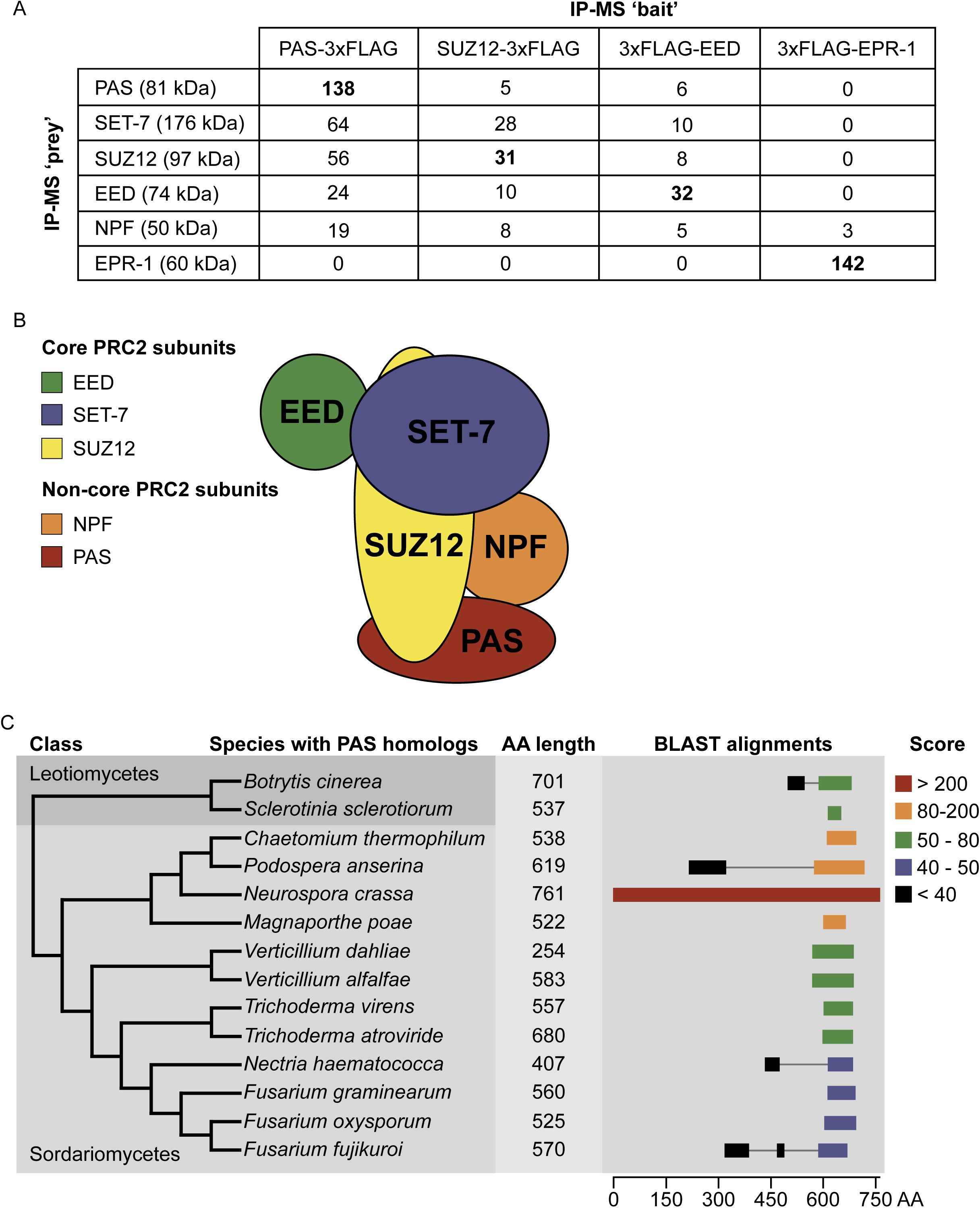
PAS is a **<u>P</u>**RC2 **<u>a</u>**ccessory **<u>s</u>**ubunit with homologs in other fungi. (A) Exclusive unique peptide counts from immunoprecipitation followed by mass spectrometry (IP-MS) experiments. Each column represents the results of a single IP-MS with the indicated ‘baits.’ The 3xFLAG-EED dataset is previously published (10). The 3xFLAG-EPR-1 sample is a negative control. (B) Diagram of putative PRC2 complex containing PAS. Adapted from (30). (C) Phylogenetic tree highlighting representative fungal species in Sordariomycetes and Leotiomycetes that have predicted PAS homologs. Length, in amino acids (AA), of predicted PAS homologs in representative species is indicated to the right of species names. Results of the <u>B</u>asic <u>L</u>ocal <u>A</u>lignment <u>S</u>earch <u>T</u>ool (BLAST), using *Neurospora crassa* PAS as the query, for each of the indicated species, is displayed to the right of homolog length. Solid colored bars indicate an aligned region. Grey lines are for visual reference. Amino acid position in PAS is indicated below the alignments. Alignment scores are color-coded according to the legend.

### PAS homologs are present in a variety of fungal lineages

In an effort to determine if homologs of PAS exist outside of *N. crassa*, we performed an iterative sequence similarity search (19). We detected homologs of PAS in lineages of Sordariomycetes and Leotiomycetes (Fig. 5C). The majority of these fungal species either have H3K27 methylation experimentally validated or inferred from their genomic sequence (20), supporting the notion that the detected PAS homologs associate with PRC2. Sequence alignments of the detected PAS homologs from representative species compared to *N. crassa* PAS revealed that significant sequence similarity between the proteins is restricted to the C-terminus of *N. crassa* PAS (Fig. 5C). In addition, the total predicted length of PAS homologs is variable (Fig. 5C), further highlighting the divergence of these homologs. We conclude that while apparent PAS homologs are in lineages of Sordariomycetes and Leotiomycetes, they exhibit considerable sequence divergence.

## DISCUSSION

Ever since the identification of Polycomb response elements, sequences that reliably induce H3K27 methylation in *Drosophila melanogaster* (21), there have been searches for *cis*- and *trans*-acting factors that direct PRC2 activity in other organisms. In plants, this line of inquiry has been fruitful, with the identification of sequence-specific transcription factors that directly recruit PRC2 (22–24). In mammals, unmethylated CpG islands can recruit PRC2 (25–27), but the mechanism of this recruitment is unclear. Recently, non-core subunits of mammalian PRC2.1 and PRC2.2 were demonstrated to be required to target PRC2 (28, 29), but how these structurally diverse components target specific genomic sites is unknown. Defining the full complement of *cis*- and *trans*-acting factors that recruit PRC2 and understanding their mechanism of action remains an outstanding challenge.

Utilizing a relatively simple eukaryote, the fungus *N. crassa*, we performed a forward genetic selection to identify novel factors implicated in Polycomb repression. This yielded an unknown accessory component of *N. crassa* PRC2, PAS (NCU04278), that is necessary for subtelomeric H3K27 methylation and associated gene silencing. Interestingly, the H3K27 methylation defect observed in Δ*pas* strains bears a striking resemblance to strains lacking the PRC2 component NPF (RBBP4/7 in mammals; p55 in *Drosophila*) (Fig. 3A), although their respective defects are distinct (Fig. 4A). Homologs of NPF are known to associate with other protein complexes (11); therefore, the observed H3K27 methylation defects in Δ*npf* strains could in principle be due to non-PRC2 activities of NPF. However, the finding that PAS results in H3K27 methylation defects similar to Δ*npf* strains, but appears to only interact with PRC2 components, suggests that the H3K27 methylation defects in Δ*npf* strains are largely due to the dysfunction of PRC2. Considering that the *in vivo* role of mammalian NPF homologs (RBBP4/7) in PRC2 function cannot be tested since they are essential for proliferation (30), studies from more amenable organisms, such as *N. crassa*, are extremely valuable.

Our research group has previously identified telomere repeats, (TTAGGG)n, as effective inducers of H3K27 methylation at chromosome ends in *N. crassa* (17). This subtelomeric H3K27 methylation is largely lost in Δ*pas*, Δ*npf* and Δ*tert* (*te*lomerase *r*everse *t*ranscriptase) strains alike (Fig. 3A) (17). However, the mechanism connecting these *cis*- and *trans*-acting factors is unclear. In mammals, the accessory subunits of PRC2 are known to affect both the catalytic activity and the recruitment of PRC2 to chromatin (28–30). Conceivably, PAS and NPF may work together to stimulate PRC2 activity on ‘non-ideal’ subtelomeric targets and play no role in recruitment. It is also possible, however, that they recruit PRC2 specifically to subtelomeres without influencing catalysis. Both activity and recruitment models can equally account for the observed losses and gains of H3K27 methylation in Δ*pas* strains.

The location of the mutations in the identified *pas^UV^* alleles and the sequence alignments of predicted PAS homologs demonstrate that the C-terminus of PAS is a critical, conserved region. This invariant C-terminal region may be responsible for assembly of PAS into PRC2, whereas the divergent N-termini may dictate chromatin targeting or regulation of PRC2 activity that is species-specific. Although *N. crassa* PAS is responsible for subtelomeric H3K27 methylation, homologs of PAS in other species may be responsible for H3K27 methylation present at diverse genomic regions. Future work in other fungal species should elucidate the conservation of function, or lack thereof, of PAS-containing PRC2 complexes.

## MATERIALS AND METHODS

### Strains, media and growth conditions

All *N. crassa* strains used in this study are listed in Table S1 (supplemental material). Preparation of media and growth conditions for experiments were carried out as previously described (12).

### Selection for mutants defective in H3K27 methylation-mediated silencing

Conidia from strain N6279 were mutagenized with ultraviolet light and challenged with Hygromycin B and Nourseothricin to select for antibiotic-resistant colonies as previously described (12). Primary mutants were rendered homokaryotic by crossing to strain N3756.

### Whole genome sequencing, mapping and identification of *pas^UV1,2^* alleles

This was performed as previously described (12). Briefly, antibiotic-resistant, homokaryotic mutants were crossed to a “Mauriceville” wild-type strain (13) and antibiotic-resistant progeny were pooled for whole genome sequencing. Mapping of the critical mutations was performed as previously described (12, 14). FreeBayes and VCFtools were used to identify novel genetic variants present in our pooled mutant genomic DNA (31, 32).

### RNA isolation, RT-qPCR, and mRNA-seq

Extraction of total RNA from germinated conidia was performed as previously described (12), and either used for cDNA synthesis and subsequent qPCR (see Table S5 in supplemental material for primers) (12) or mRNA-seq library preparation (16). Mapping and analysis of gene expression levels was performed as previously described (12).

### Chromatin immunoprecipitation (ChIP), ChIP-qPCR, and ChIP-seq

H3K27me2/3 ChIP using anti-H3K27me2/3 antibody (Active Motif, 39536) was performed as previously described (12) and the isolated DNA used for qPCR (see Table S6 in supplemental material for primers) or prepared for sequencing (12). Mapping, visualization, and analysis of H3K27me2/3 ChIP-sequencing reads was performed as previously described (12). Spearman’s correlation coefficient analysis was performed using deepTools2 (33) on the Galaxy public server (34).

### Western blotting

*N. crassa* tissue lysates were prepared as previously described (12) and used for western analysis. Anti-H3K27me3 (Cell Signaling Technology, 9733) and anti-hH3 (Abcam, ab1791) primary antibodies were used with IRDye 680RD goat anti-rabbit secondary (LI-COR, 926-68071). Images were acquired with an Odyssey Fc Imaging System (LI-COR) and analyzed with Image Studio software (LI-COR).

### Immunoprecipitation followed by mass spectrometry (IP-MS)

Strain N7807 (over-expressing *NCU04278*::C-Gly::3xFLAG) or strain N4666 (endogenous *suz12*::C-Gly::3xFLAG) was grown for 7-10 days in a 250 mL flask containing 50 mL of Vogel’s minimal medium containing 1.5% sucrose and 1.5% agar. Approximately 1×10^9^ conidia were collected from each culture and filtered through sterile cheesecloth. Filtered conidia were used to inoculate 1 L of Vogel’s minimal medium containing 1.5% sucrose and then grown for 16 hours shaking (150 RPM) at 30 °C. Tissue was collected by filtration using a Buchner funnel and washed with water. Dry tissue (∼10 g) was ground using a 6870 Freezer/Mill Cryogenic Grinder (SPEXSamplePrep). Ground tissue was added to 40 mL Extraction Buffer (EB) (50 mM HEPES [pH=7.5], 150 mM NaCl, 10 mM EDTA, 10% glycerol, 0.02% NP-40, 1X Halt Protease Inhibitor Cocktail (Thermo Scientific, 78438)) to achieve a final volume of 50 mL, and then suspended by rotation at 4 °C for 1.5 hours. Insoluble material was pelleted with one 10-minute centrifugation step at 2,000 RPM and two consecutive 10-minute centrifugation steps at 8,000 RPM. Soluble material was pre-cleared with 250 µL of equilibrated Protein A agarose (Invitrogen, 15918014) with rotation for 1 hour at 4 °C. The Protein A agarose was pelleted by centrifugation at 2,000 RPM and the supernatant collected, which was then incubated with 400 µL of equilibrated ANTI-FLAG M2 affinity gel (Sigma-Aldrich, A2220) overnight, rotating at 4 °C. Resin was pelleted by centrifugation at 1,000 RPM and supernatant removed. Resin was washed with EB, rotated at 4 °C for 10 minutes, and pelleted with a 1,000 RPM spin five consecutive times. All liquid was removed after the final spin. Protein was eluted twice from the resin by incubating with 300 µL of 500 µg/mL 3X Flag Peptide (APExBIO, A6001) rotating at 4 °C for 20 minutes, and one final wash with 300 µL of EB. Eluate was precipitated with trichloroacetic acid (10% final concentration) on ice for 1 hour, pelleted by centrifugation at 14,000 RPM, and washed three times with ice-cold acetone. Pellet was air-dried by placing in heat block at 100 °C for 30 seconds. Samples were sent to and processed by the UC Davis Proteomics Core Facility for mass spectrometry and subsequent analysis.

### Bioinformatic analysis of PAS homologs

Homologs of *N. crassa* PAS (accession number: Q1K790) were detected using JACKHMMER iterative search (19), which converged after three consecutive iterations. Regions of significant alignment between detected PAS homologs from representative species and *N. crassa* PAS were analyzed using the BLAST web server (35). Phylogenetic relationships between fungal species were based on previous work (36).

### Replacement of *NCU04278* with *trpC::nat-1*

The 5’ and 3’ flanks of *NCU04278* were PCR-amplified from wild-type genomic DNA with primer pairs 6405 and 6406 (5’) and 6409 and 6410 (3’) (see Table S3 in supplemental material). The 5’ and 3’ flanks were separately PCR-stitched to plasmid 3237 (source of *trpC::nat-1*) using primer pairs 6401 and 4883, and 4882 and 6351, respectively. These two ‘split-marker’ PCR products were co-transformed into strain N2930 and *NCU04278* gene replacements were selected on Nourseothricin-containing medium.

### Generation of an *N. crassa* strain expressing PAS::3xFLAG

*NCU04278* was PCR-amplified from wild-type genomic DNA with primers 6397 and 6398 (see Table S4 in supplemental material), and cloned into plasmid 2401 (37) using XbaI and PacI restriction sites to create plasmid 3340 (see Table S2 in supplemental material). Plasmid 3340 was subjected to Sanger sequencing using primers 6397, 6407, 6426, 6427, and 6715 to verify that the sequence matched that of wild type. Plasmid 3340 was linearized with NdeI and targeted to *his-3* in N6762, as previously described (38). A *his-3^+^* primary transformant was then crossed to N7742 to generate N7807.

### Data availability

Whole-genome sequencing data from *pas^UV1^* and *pas^UV2^* mapping experiments is available from the Sequence Read Archive (PRJNA559544). The results of our mRNA-seq and ChIP-seq experiments are available on the NCBI GEO database (GSE140787). Results of the mass spectrometry experiments are included in the supplemental material.

## SUPPLEMENTAL MATERIAL

Supplemental material for this article may be found at …

## ACKNOWLEDGMENTS

We wish to thank J. Lyle and A. Leiferman for help in genetic mapping of UV-generated mutants, and M. Salemi at the UC Davis Proteomics Core Facility for the mass spectrometry work. This study was funded by the National Institute of General Medical Sciences (GM127142 and GM093061 to E.U.S.) and the American Heart Association (14POST20450071 to E.T.W.). K.J.M. was supported in part by the National Institutes of Health (T32-HD007348).

---

**Table S1.**
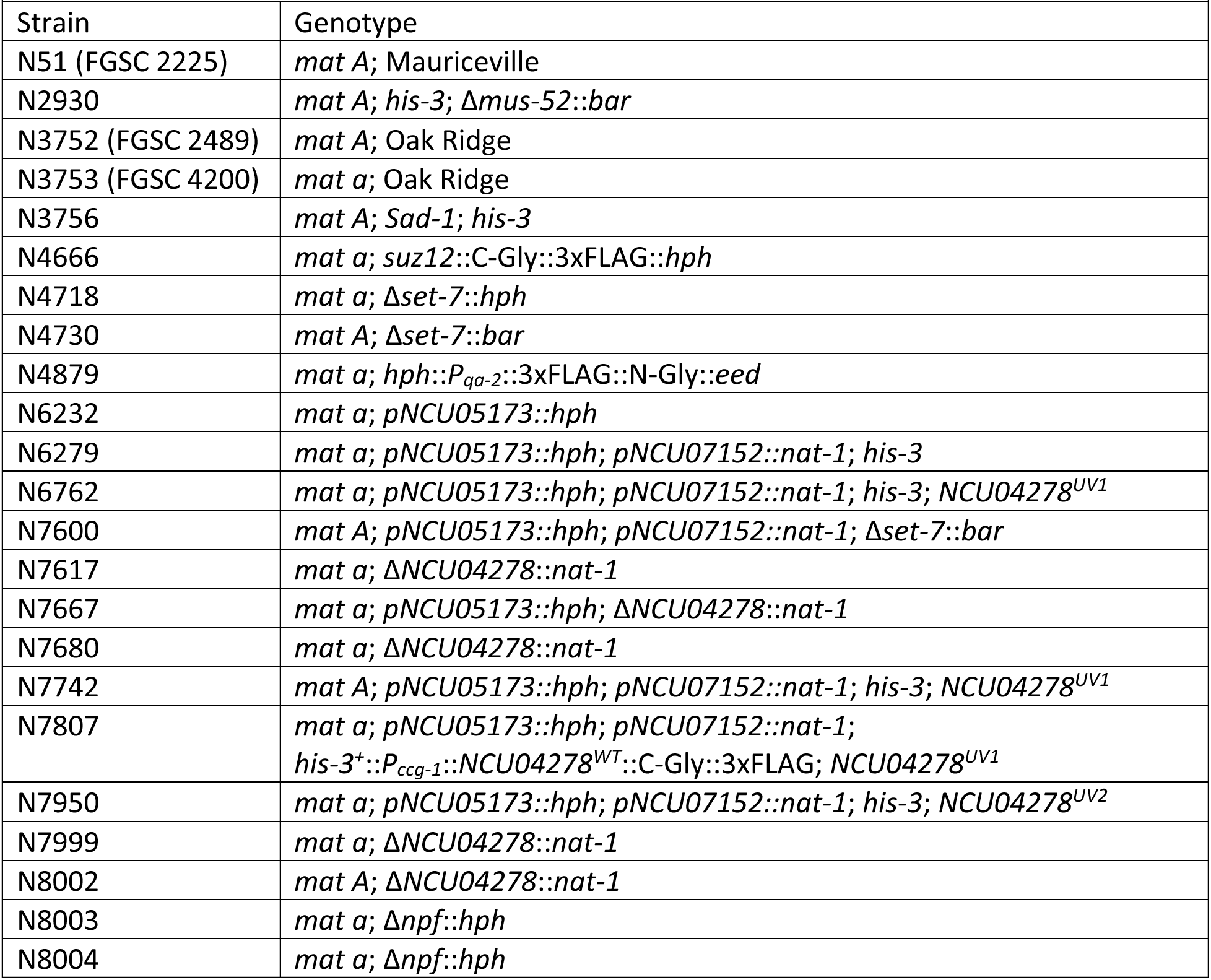
*N. crassa* strains

**Table S2.** Plasmids

| Table S2. Plasmids |  |
| --- | --- |
| Plasmid | Description |
| 2401 | *P<sub>CCG-1</sub>::C-Gly::3xFLAG* |
| 3237 (FGSC 10598) | pAL12 – source of *nat-1* |
| 3340 | *P<sub>CCG-1</sub>::NCU04278<sup>WT</sup>::C-Gly::3xFLAG* |

**Table S3.**
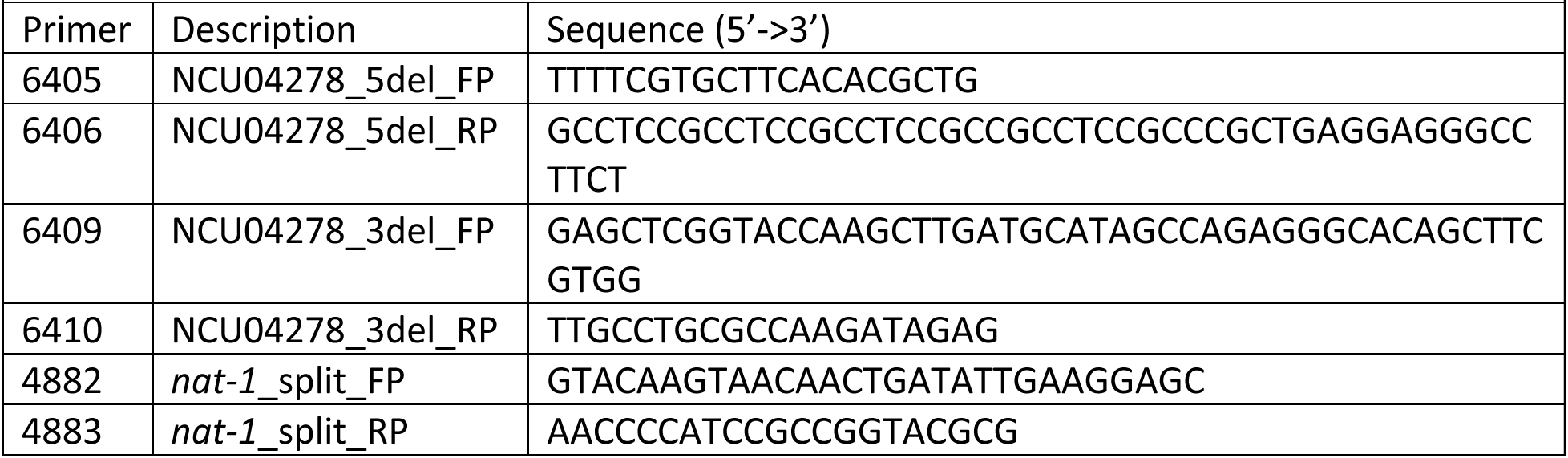
Primers for replacement of *NCU04278* with *trpC::nat-1*

**Table S4.**
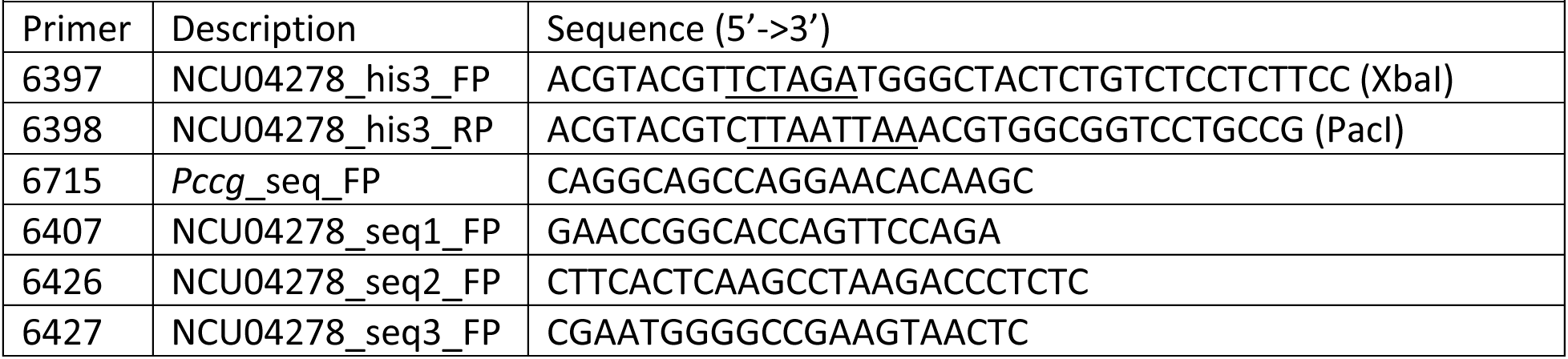
Primers for generation and verification of *his-3^+^::P_CCG-1_::NCU04278*::3xFLAG construct

**Table S5.**
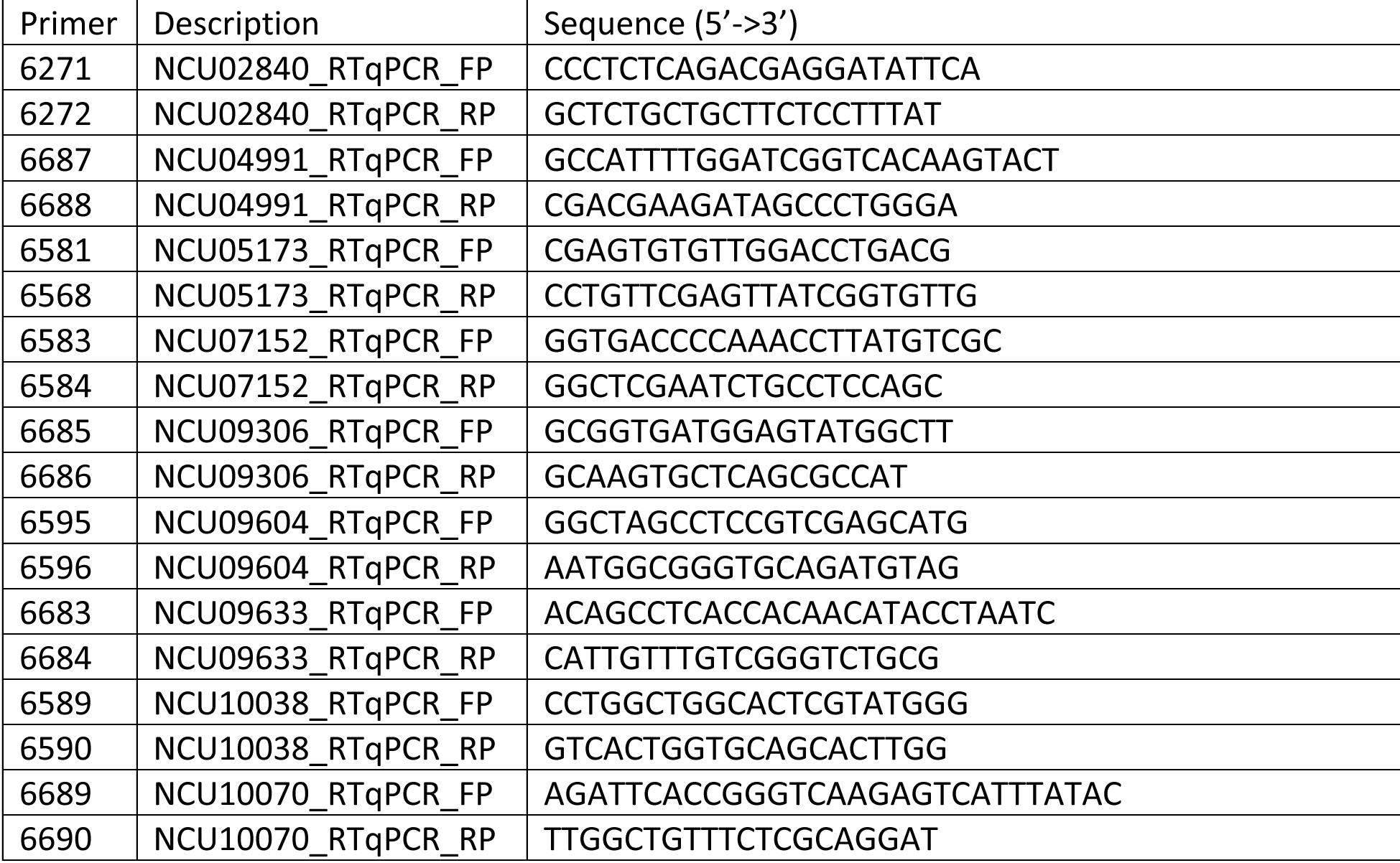
RT-qPCR primers

**Table S6.**
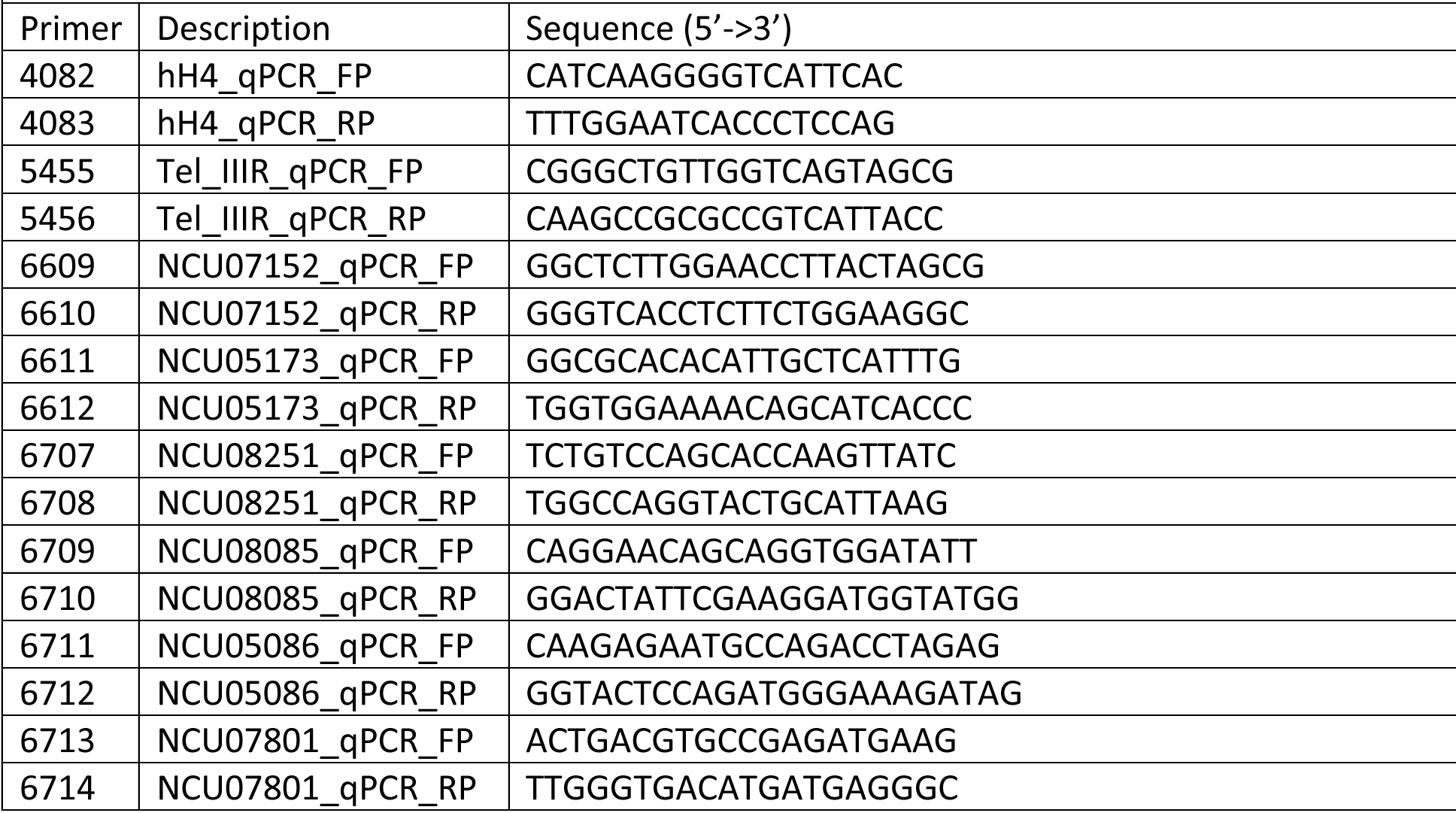
ChIP-qPCR primers

